# Tumor-Associated Macrophages in Meningiomas: An Independent Prognostic Factor for Poor Survival Outperforming the Benefits of T cells

**DOI:** 10.1101/2024.12.11.627726

**Authors:** Catharina Lotsch, Fang Liu, Rolf Warta, Gerhard Jungwirth, Carmen Rommel, Mandy Barthel, Katrin Lamszus, Almuth F. Kessler, Niels Grabe, Mario Loehr, Ralf Ketter, Christian Senft, Manfred Westphal, Felix Sahm, Sandro Krieg, Andreas Unterberg, Matthias Simon, Andreas von Deimling, Christel Herold-Mende

**Author notes:** Authors contributed equally to this work (Catharina Lotsch, Fang Liu, Rolf Warta, Gerhard Jungwirth). **Address correspondence to:** Prof. Dr. rer. nat. Christel Herold-Mende Division of Experimental Neurosurgery, Department of Neurosurgery, University of Heidelberg, INF400, 69120 Heidelberg, Germany.

## Abstract

**Background:** Tumor-associated macrophages (TAMs) represent the main immune cell population in various brain malignancies. To elucidate their biological impact in the tumor microenvironment (TME) of meningiomas (MGMs), we assessed TAM numbers, activation state, malignancy- and survival-associated changes, as well as their association with tumor-infiltrating T lymphocytes (TILs).

**Methods:** TAM infiltration was analyzed in a multicenter cohort of 195 clinically well-annotated cases (follow-up >5 years, *n*=120 newly-diagnosed and *n*=75 recurrent MGMs) enriched for higher-grade MGMs. TAMs and M2-TAMs were quantified by tissue cytometry on whole-tumor sections. Further, we assessed levels of 27 cyto- and chemokines in a subset of tissues (*n*=46 cases), and re-analyzed our previously published T cell infiltration (*n*=94 cases) and expanded microarray (*n*=97 cases) datasets.

**Results:** Newly-diagnosed MGMs showed a substantial but highly heterogeneous TAM infiltration that was four times higher than for TILs. Anti-inflammatory M2-TAMs were increased in higher WHO grade tumors and in recurrent MGMs. Importantly, high M2-TAM infiltration was associated with poor progression-free survival independent of other prognostic confounders and even mitigated the beneficial prognostic effect of TIL infiltration. Additional cytokine, gene expression and pathway analyses corroborated the presence of an immunosuppressive niche in M2-TAM-enriched MGMs.

**Conclusions:** Altogether, higher numbers of TAMs and M2-TAMs appear to be a hallmark of clinically aggressive behavior in newly-diagnosed and recurrent MGMs. Unlike TILs, immunosuppressive TAMs seem to play a dominant negative role in the immunological landscape of MGMs, highlighting M2-TAMs to be an attractive treatment target for immunotherapeutic approaches.

**Translational Relevance of the Study:** Meningiomas (MGMs) are typically regarded as benign neoplasms, however there is a substantial proportion of clinically aggressive tumors that are refractory to standard treatment modalities and demand for the development of novel therapeutic approaches such as immunotherapy. This is the first comprehensive study reporting malignancy- and progression-associated changes of tumor-associated macrophages (TAMs), their polarization state, their association with tumor-infiltrating T lymphocytes (TILs), and their impact on patient survival in a large multicenter cohort of 195 tumors containing substantial numbers of clinically aggressive cases. Notably, we identified higher numbers of immunosuppressive M2-TAMs as an independent prognostic factor for poor survival, overriding the beneficial prognostic effects of TILs. Thus, our data highlight an important role of immunosuppressive M2-TAMs on tumor malignancy and progression, and further suggest targeting macrophages as a treatment strategy to improve the success of immunotherapeutic approaches in MGMs.

**Key points:**

- Meningiomas are highly infiltrated by immunosuppressive M2-TAMs.
- High M2-TAM numbers are an independent negative prognostic factor for patient survival.
- High TAM infiltration mitigates the beneficial prognostic impact of TILs.

## INTRODUCTION

Meningiomas (MGMs) are the most common primary intracranial tumors in adults, representing over a third of all the neoplasms in the CNS.^1–3^ Although the majority of MGMs are slow-growing and histologically classified as benign (WHO°1), there is a substantial subset of tumors exhibiting aggressive clinical behavior, resulting in recurrence rates at 5 years of 10-15% for WHO°1, 50% for WHO°2 and 90% for WHO°3 MGMs, respectively.^4,5^ Standard treatment options are limited to surgery and radiotherapy, and despite intensive research efforts no systemic treatment options are yet available for clinically aggressive tumors.^6,7^ With the breakthrough of immune checkpoint blockade (ICB) therapy in various other cancer types, research into the immunobiology of MGMs has received a great surge of interest, but there is still insufficient knowledge about the tumor’s immune microenvironment and its impact on disease progression and patient outcome.^8^

In comparison to other malignancies, MGMs are characterized by a low tumor mutational burden, lower lymphocyte numbers and a predominantly immunosuppressive micromilieu, particularly in higher-grade tumors.^9,10^ First studies have shed light into the complex immunological landscape in MGMs and reported also the infiltration of immunosuppressive cells of myeloid origin including myeloid-derived suppressor cells (MDSCs)^11,12^ and tumor-associated macrophages (TAMs).^13,14^ Together these studies provided first evidence that meningiomas seem to be largely infiltrated by TAMs. However, until now, an integrative analysis of the phenotype and functional role of TAMs with regard to T cell prevalence, tumor behavior and patient survival in a large and well-balanced tumor cohort is missing for this cancer entity.

In general, TAMs are recognized as highly heterogeneous and plastic cells with both their phenotype and function being strongly influenced by local microenvironmental cues in the tumor milieu.^15^ TAMs are regularly dichotomized into M1/M2 macrophage polarization classes, in which M1-TAMs are considered as anti-tumoral while M2-TAMs are presented as pro-tumoral.^16,17^ M2-TAMs are characterized by increased expression of immune checkpoint ligands (PD-L1), secretion of anti-inflammatory cytokines (IL-10, TGF-β), and upregulated expression of scavenger receptors (CD163, CD204, CD206).^15,18,19^ In various cancers, immunosuppressive M2-TAMs play an important role in tumorigenesis and disease progression and their presence is generally associated with a poor prognosis.^15,16,20^

To shed more light on the role of TAMs in MGMs and especially their impact on tumor behavior and patient survival, in this study we report TAM frequencies in a large cohort of almost 200 clinically well-annotated MGM cases, enriched for clinically aggressive tumors. We found a malignancy- and progression-increased TAM and M2-TAM infiltration. Notably, in our multivariate analysis we identified high TAM infiltration as an independent prognostic factor for inferior progression-free survival (PFS) in patients with newly-diagnosed MGMs, which even outperformed the beneficial impact of higher numbers of tumor-infiltrating T lymphocytes (TILs) in the same study sample. Taken together, our data suggest that high TAM and M2-TAM infiltration seems to be a hallmark of clinically more aggressive behavior in MGMs, defining M2-TAMs as an attractive treatment target for cancer immunotherapy in MGM patients in the future.

## MATERIALS AND METHODS

### Samples and patient characteristics

A total of 195 meningioma specimens (WHO°1 *n*=43; WHO°2 *n*=97; WHO°3 *n*=62) were obtained from female and male patients undergoing surgical resection in the Departments of Neurosurgery at University Hospitals Heidelberg, Bonn, Hamburg, Homburg, Frankfurt, and Würzburg, Germany as part of the “FORAMEN” and “KAM” consortia.^21–23^ The use of tissue samples was approved by institutional review boards at each institute in accordance with the Declaration of Helsinki. Written informed consent was obtained from all patients. Tumor specimens were immediately snap-frozen after surgery and stored at −80°C until further processing. Tumor cell content ≥60% was confirmed for all samples by an experienced neuropathologist (AvD). Tumor specimens with a tumor cell content <60% were excluded. Clinical data were collected using a detailed questionnaire and are summarized in Table 1.

**Table 1:** Clinicopathological characteristics of patients with newly-diagnosed and recurrent meningioma.

| Variable | Newly-diagnosed MGMs (n=120) |  |  | Recurrent MGMs (n=75) |  |  |
| --- | --- | --- | --- | --- | --- | --- |
|  | n | Patients (%) | Median (range) | n | Patients (%) | Median (range) |
| Sex |  |  |  |  |  |  |
| Male | 46 | 38.33 |  | 41 | 54.66 |  |
| Female | 74 | 61.66 |  | 34 | 45.33 |  |
| Age at 1<sup>st</sup> diagnosis (years) |  |  | 60.8(24.0-87.6) |  |  | 55.0 (18.0-86.5) |
| WHO grade |  |  |  |  |  |  |
| WHO°1 | 33 | 27.50 |  | 10 | 13.33 |  |
| WHO°2 | 63 | 52.50 |  | 32 | 42.66 |  |
| WHO°3 | 24 | 20.00 |  | 33 | 44.00 |  |
| Subtype |  |  |  |  |  |  |
| Transitional | 12 | 10.00 |  | 9 | 12.00 |  |
| Fibroblastic | 8 | 6.66 |  |  |  |  |
| Meningothelial | 8 | 6.66 |  |  |  |  |
| Angiomatous | 1 | 0.83 |  |  |  |  |
| Secretory | 1 | 0.83 |  |  |  |  |
| Atypical | 52 | 43.33 |  | 25 | 33.33 |  |
| Anaplastic | 12 | 10.00 |  | 28 | 37.33 |  |
| Rhabdoid | 1 | 0.83 |  | 1 | 1.33 |  |
| Papillary | 2 | 1.66 |  |  |  |  |
| NA | 23 | 19.16 |  | 12 | 16.00 |  |
| Localization |  |  |  |  |  |  |
| Convexity | 50 | 41.66 |  | 21 | 28.00 |  |
| Cranial base | 29 | 24.16 |  | 16 | 21.33 |  |
| Falx | 14 | 11.66 |  | 15 | 20.00 |  |
| Parasagittal | 17 | 14.16 |  | 13 | 17.33 |  |
| Tentorial | 4 | 3.33 |  | 4 | 5.33 |  |
| Other multiple or NA | 6 | 5.00 |  | 6 | 8.00 |  |
| Resection grade |  |  |  |  |  |  |
| Simpson 1 | 67 | 55.83 |  | 29 | 38.66 |  |
| Simpson 2 | 32 | 26.66 |  | 20 | 26.66 |  |
| Simpson 3 | 16 | 13.33 |  | 12 | 16.00 |  |
| Simpson 4 | 4 | 3.33 |  | 12 | 16.00 |  |
| Simpson 5 | 0 | 0.00 |  | 1 | 1.33 |  |
| NA | 1 | 0.83 |  | 1 | 1.33 |  |
| Postoperative radiotherapy |  |  |  |  |  |  |
| Yes | 34 | 28.33 |  | 32 | 42.66 |  |
| No | 81 | 67.50 |  | 41 | 54.66 |  |
| NA | 5 | 4.16 |  | 2 | 2.66 |  |
| Postoperative chemotherapy |  |  |  |  |  |  |
| Yes | 0 | 0.00 |  | 7 | 9.33 |  |
| No | 117 | 97.50 |  | 64 | 85.33 |  |
| NA | 3 | 2.50 |  | 4 | 5.33 |  |

Abbreviations: n, number; NA, not available; MGM, meningioma

### Multicolor immunofluorescence staining

Multicolor immunofluorescence staining was performed on acetone-fixed cryosections (5-7μm). To quantify TAM and M2-TAM subpopulations, a combination of primary antibodies specific for CD68 (mouse, 1:25, Agilent Cat# M0718, RRID:AB_2687454), CD163 (mouse, 1:300, MCA1853, Bio-Rad Cat# MCA1853T, RRID:AB_2074539), and CD204/MSR1 (rabbit, 1:200, Sigma-Aldrich Cat# HPA000272, RRID:AB_1846269) were used as described previously.^24^ Briefly, CD163 and CD204 primary antibodies were diluted with Antibody Diluent (Dako). For CD68 staining, primary antibody was coupled to AlexaFluor488 with Zenon labelling kit according to the manufacturer’s instructions (Thermo Fisher Scientific). Secondary antibodies were used as follows: anti-mouse AlexaFluor647 (1:200, Thermo Fisher Scientific), anti-rabbit AlexaFluor555 (1:800, Invitrogen) for staining of CD163 and CD204. Secondary antibodies were diluted with DPBS-containing DAPI (Thermo Fisher Scientific) at 1:1,000 to stain nuclei. Primary anti-CD163, anti-CD204 and secondary antibodies were incubated for 1h, anti-CD68 primary antibody was incubated for only 20min. Human tonsil tissue and isotype-matched antibodies (rabbit IgG, x0936, Dako; rat IgG2b, 14-4031, eBioscience; and mouse IgG1, ab91353, Abcam) served as positive and negative controls, respectively.

### Tissue cytometry-based image analysis

Image analysis was performed in a semiautomated set-up at a single-cell level with subsequent phenotypic hierarchical clustering as described before.^22,24^ In brief, high-resolution automated multiple image alignments of whole-tissue sections were acquired using a 20x objective on an Olympus IX51 microscope equipped with a XM10 Camera (Olympus). The Olympus cellSens Dimension Software (version 1.9) was used for image acquisition. Automatic detection and context-based quantification of TAM infiltration by immunofluorescence markers was performed by the StrataQuest Software (version 5.0.1, TissueGnostics GmbH). Regions of interest (ROI) were manually defined depending on histology and quality of the section to exclude adjacent normal brain or necrotic areas. ROIs were drawn in the slide overview using software-based mark-up tools. Quantification was solely performed in areas with high tumor cell content (≥60%).

Automatically detected cells were visualized in scattergrams and gated according to defined gating schemes for the expression of nucleic and cell surface markers (Supp. Fig. S1). Cutoff between positive- and negative-gated cells was validated by backward gating. To enable robust and reliable cell quantification, strict parameters by means of nuclear size, staining intensity, and background threshold were defined. Cell nuclei were detected based on DAPI staining and used as origin to generate a growing mask over the cytoplasm to the cell membrane. Based on this mask, TAMs were analyzed regarding cell surface expression of CD68 and DAPI (Suppl. Fig. S1). M2-TAMs were defined by the cell surface expression of CD68 and CD163 and/or CD204. For statistical analysis, number of cells was given in percent of total cell count (%TCC, defined as total number of DAPI^+^ nuclei without further distinction of cell types).

### Luminex assay

Luminex analysis was performed using the Bio-Plex Pro Human Cytokine 27-plex Assay (Bio-Rad) according to manufacturer’s protocol. In short, protein was isolated from a total of 46 MGM specimens (newly-diagnosed MGMs: WHO°1 *n*=7, WHO°2 *n*=19, WHO°3 *n*=6; recurrent MGMs: WHO°1 *n*=2, WHO°2 *n*=9, WHO°3 *n*=3) using the Bio-Plex Cell Lysis Kit (Bio-Rad) according to the manufacturer’s instructions. The protein concentrations were determined by Pierce BCA Protein Assay Kit (Thermo Fisher Scientific) and subsequently, all lysates were diluted to 1mg/mL. In 96-well assay plate, 50µL of 1x beads were added and then washed with 100µL wash buffer twice. Thereafter, 50µL of standards, samples and controls were added and incubated on a shaker at 850rpm for 30min, followed by a washing step. Then, 50µL 1x streptavidin-PE were added to each well and the plate was incubated at 850rpm for 10min. Thereafter, all wells were washed with 3×100µL. Finally, the beads were resuspended in 125µL assay buffer, shaked at 850rpm for 30s. Data acquisition and analysis was done using the Luminex 100 Bio-Plex System and the Bio-Plex Manager Software version 6.1 (Bio-Rad).

### Microarray analysis

For microarray analysis, we re-used our previously published microarray dataset GSE74385 (*n*=62 cases)^21^ and extended it with additional *n*=35 MGM cases. As described previously, total RNA was extracted from MGM tissue specimens using the AllPrep DNA/RNA/Protein Kit (Qiagen) according to the manufacturer’s instructions. RNA concentration and integrity were analyzed using the 2100 Bioanalyzer (Agilent). For microarray analysis, 1µg total RNA of each tumor specimen was subjected to the Genomics Core Facilities of the German Cancer Research Center (DKFZ, Heidelberg, Germany). After quality control, purification and cDNA synthesis, samples were hybridized to Human HT-12 V.4.0 BeadChip arrays (Illumina) according to the manufacturer’s instructions. Raw-intensity data of the microarrays were further processed and normalized as described before using R programming [www.r-project.org].^21^ The data were analyzed using the following packages: vsn, limma, msigdbr, clusterprofiler, and enrichplot. Following vsn normalization, differential expression was determined by limma using a model that included TAM and TIL infiltration grouping, sex, age at diagnosis, histology, and newly-diagnosed or recurrent status. Gene Ontology (GO) term and Reactome gene set enrichment analysis (GSEA) was conducted using the Molecular Signatures Database (MSigDB) collections C5 (GO) and C2 (REACTOME). The level of significance for GSEA was set to adjusted *P*-value (*P*_adj_) <0.05.

### Statistical analysis

Data were analyzed by R (Version 4.4.1, survival package) or GraphPad (Version 9.0.0). Differences between two groups (WHO°1 vs. °2; WHO°2 vs. °3; WHO°1 vs. °3; newly-diagnosed vs. recurrent MGMs; high vs. low infiltration divided by the median) were calculated using Mann–Whitney-U tests for non-parametric data or Student’s unpaired t tests for parametric data. Data are presented with median values for non-parametric data and with mean values for parametric data. Kaplan–Meier plots were used to visualize survival estimates, whereas comparison of survival differences was done by log-rank test and Cox proportional hazard (PH) models. Progression-free survival (PFS) analyses were only performed on patients with Simpson°I-III resection with a follow-up time of at least 60 months, and were otherwise excluded from the survival analysis. Variables reaching significance in univariate analyses were further included in a multivariate model to assess the independence of clinical covariates. *P*-values <0.05 were considered significant: *, *P*<0.05; **, *P*<0.01, ***, *P*<0.001, ****, *P*<0.0001).

## RESULTS

### Malignancy- and progression-related increase of TAMs and M2-TAMs in meningioma

To study the infiltration of TAMs and their polarization in MGMs, we performed multicolor immunofluorescence staining with antibodies against CD68, CD163, and CD204 in a cohort of 195 clinically well-annotated cases including *n*=120 newly-diagnosed and *n*=75 recurrent MGMs (Fig. 1A, Table 1). For newly-diagnosed MGMs, the study cohort consisted of 33 WHO°1, 63 WHO°2, and 24 WHO°3 tumors. The median age of patients was 60.8 years at the time of the first diagnosis with a female to male ratio of 1.6 to 1.0. To assess whether tumor recurrence results in altered TAM infiltration, we analyzed a second cohort of 75 recurrent MGMs comprising a substantial number of high-grade tumors (WHO°1, *n*=10; WHO°2, *n*=32; WHO°3, *n*=33). To increase reliability, we analyzed whole tissue sections of newly-diagnosed (median area: 11.11mm², range: 1.94-36.25mm²) and recurrent MGMs (median area: 10.7mm², range: 0.97-31.15mm²) by tissue cytometry-based image analysis (Suppl. Fig. S1). We found TAM infiltration, assessed by the number of TAMs (CD68^+^) relative to the total cell count (TCC) within the tumor section, to be highly heterogeneous across MGM specimens: TAM percentages varied widely from 0.01% to 31.8% with a median of 2.47% CD68^+^/TCC for newly-diagnosed MGMs (Fig. 1B), and shifted towards a higher abundance in recurrent MGMs with a median of 3.22% CD68^+^/TCC (Fig. 1C). Median TAM infiltration for newly-diagnosed MGMs increased with WHO grade: from 1.9% in grade 1 to 2.4% in grade 2 (increase of 26% from grade 1), and to 4.1% in grade 3 tumors (increase of 116% from grade 1), but without reaching a level of significance (Fig. 1D; WHO°1 to °2, *P*=0.148; WHO°1 to °3, *P*=0.082; Mann-Whitney-U test).

**Figure 1:**
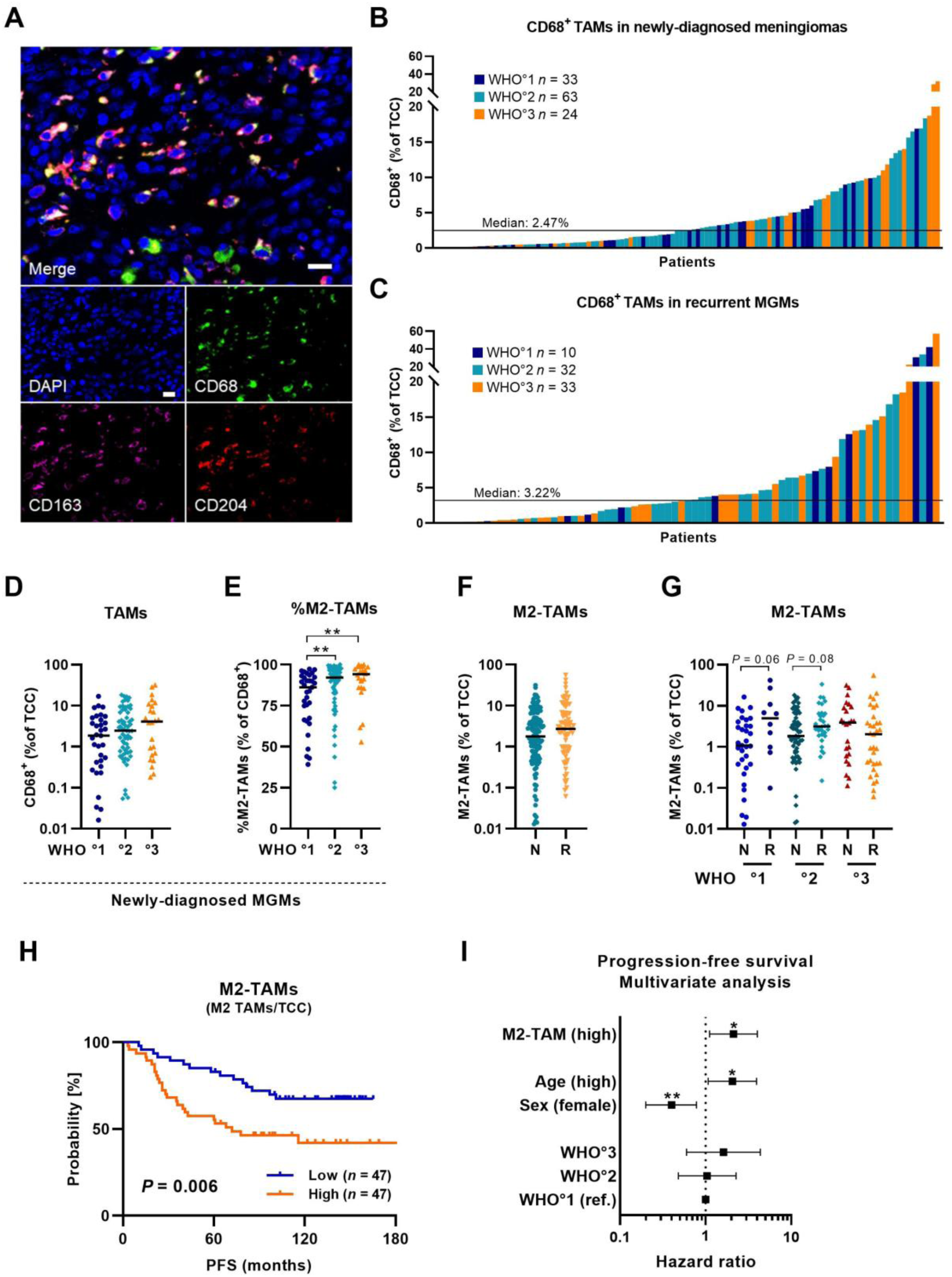
TAM infiltration in newly-diagnosed and recurrent meningiomas. **(A)** Representative fluorescent images of TAM staining in MGM tissues. Nuclei stained with DAPI in blue, TAM and M2-TAM stained with CD68 in green, CD163 in purple, CD204 in red. Scale bar: 20µm. **(B)** TAM infiltration (CD68^+^/TCC) in newly-diagnosed MGMs. **(C)** TAM infiltration in recurrent MGMs. **(D)** TAM infiltration across WHO grades in newly-diagnosed MGMs. **(E)** Proportion of M2-TAMs (M2-TAMs/CD68^+^) across WHO grades in newly-diagnosed MGMs. **(F)** M2-TAM infiltration (M2-TAMs/TCC) in newly-diagnosed (N) and recurrent (R) MGMs. **(G)** M2-TAM infiltration across WHO grades in newly-diagnosed (N) and recurrent (R) MGMs. **(H)** Kaplan-Meier plot for PFS based on high (orange curve) and low (blue curve) M2-TAM infiltration in newly-diagnosed MGMs **(I)** Multivariate survival analysis for PFS including prognostic confounders (age, sex, WHO grade) and M2-TAM infiltration. Statistical significance was calculated using Mann-Whitney-U test in (D-G), log-rank test in (H), and Cox proportional hazard model in (I). Abbreviations: MGM, meningioma; N, newly-diagnosed; PFS, progression-free survival; R, recurrent; TAM, tumor-associated macrophage; TCC, total cell count. Statistical significance: *, *P*<0.05; **, *P*<0.01; ***, *P*<0.001.

Since TAMs can acquire both anti-tumorigenic M1 or tumor-supportive M2 phenotypes in the local tumor milieu, we next examined the polarization status of TAMs by applying CD204 and CD163 and thus two well-known M2-TAM markers.^15,18,19^ Therefore, CD68^+^ TAMs with an additional CD163^+^ and/or CD204^+^ detection were regarded as pro-tumorigenic and immunosuppressive M2-TAMs. Firstly, we were interested if the proportion of M2-TAMs (%M2-TAMs of CD68^+^) differed among WHO grades in newly-diagnosed MGMs. Here, the median percentage of the M2-TAM population significantly increased from grade 1 tumors to grade 2, and to grade 3 tumors (Fig. 1E; WHO°1 to °2, *P*=0.009; WHO°1 to °3, *P*=0.003; Mann-Whitney-U test). Next, we analyzed the infiltration of M2-TAMs (M2-TAMs/TCC) in the whole cohort and found a 1.5-fold higher prevalence of M2-TAMs in recurrent tumors with a median of 2.72% compared to newly-diagnosed MGMs (median of 1.78%; Fig. 1F; *P*=0.209; Mann-Whitney-U test). When analyzing M2-TAM infiltration upon recurrence within the same WHO grade, we observed a 3-fold increase of M2-TAM infiltration in recurrent grade 1 MGMs (Fig. 1G; *P*=0.055; Mann-Whitney-U test) and a 1.5-fold increase in recurrent grade 2 tumors (Fig. 1G; *P*=0.078; Mann-Whitney-U test). However, our analysis did not reach the level of significance probably due to the small sample sizes in the subgroups. Similar results were obtained for the proportion of total TAMs when comparing newly-diagnosed with recurrent tumors (Suppl. Fig. S2B).

In summary, analysis of TAM frequencies in newly-diagnosed and recurrent MGMs exhibited a highly heterogeneous but substantial TAM infiltration in the tumor tissue. We found significantly increased proportions of M2-TAMs in WHO°2 and °3 newly-diagnosed MGMs and increased frequencies of M2-TAMs especially in WHO°1 recurrent tumors.

### High M2-TAM infiltration is an independent prognostic factor for poor survival in meningiomas

Next, we analyzed whether TAM infiltration has an impact on patient outcome in MGMs. To avoid any bias in the survival analysis due to incomplete tumor resection, treatment-induced changes or a short follow-up, we only included patients who underwent a gross total tumor resection (Simpson°I-III), without any prior treatment, and with a minimum follow-up time of 60 months. For the analysis, the patient cohort was then divided into low and high infiltration groups according to the median of TAM and M2-TAM infiltration. For newly-diagnosed MGMs, survival analysis of the resulting patient cohort (*n*=94) revealed that high infiltration with M2-TAMs was significantly associated with inferior PFS (Fig. 1H; *P*=0.006; log-rank test). A similar observation for PFS was seen for total TAM infiltration in patients with newly-diagnosed MGMs (Suppl. Fig. S2C), whereas in recurrent MGMs, the presence of TAMs had no further impact on survival after recurrence (Suppl. Fig. S2D).

Subsequently, a multivariate analysis was conducted, incorporating age, sex, and WHO grade as relevant prognostic factors. This analysis revealed that high M2-TAM infiltration is an independent prognostic factor for poor PFS in patients with newly-diagnosed MGMs (Fig. 1I, Suppl. Table S1; HR 2.11; 95% CI 1.11-4.01; *P*=0.023; Cox PH model).

Altogether, high numbers of M2-TAMs were found to be associated with a poor PFS in patients with newly-diagnosed MGMs independent of other prognostic factors.

### Higher TAM infiltration is associated with an immunosuppressive micromilieu in meningiomas

TAMs play vital roles in the local tumor milieu by secreting various soluble factors, including chemokines, cytokines and growth factors, which can in particular influence the attraction and function of effector T cells.^15^ To explore the TAM-related cytokine and chemokine milieu in MGM, Luminex analyses of 27 different immune-related cyto-, chemokines and growth factors were performed in a subset of 46 tissue samples (WHO°1 *n*=9, WHO°2 *n*=28, WHO°3 *n*=9). In order to identify TAM infiltration-specific distinctions, we assessed protein levels based on a median split of TAM infiltration in the tissues. Three cytokines (IL-2, IL-5 and IL-15) had to be excluded from the analysis since protein levels were not detectable in most tissues. For the majority of the remaining 24 analyzed factors a tendency towards increased protein levels in tumors with high TAM infiltration was observed (Fig. 2A), suggesting that TAM numbers may predominantly influence the production of cyto-, chemokines and growth factors in the immune microenvironment of MGMs. Interestingly, in MGM tissues with high TAM infiltration we discovered a significant increase of G-CSF, Eotaxin, IL-1β, IL-1ra, and IL-4 (Fig. 2A; *P*=0.011 for G-CSF, *P*=0.023 for Eotaxin, *P*=0.010 for IL-1β, *P*=0.008 for IL-ra, *P*=0.020 for IL-4; Mann-Whitney-U test), as well as a tendency towards increased levels of IL-6 (Fig. 2A; *P*=0.064; Mann-Whitney-U test). Particularly, the cytokines IL-1β, IL-1ra, IL-4, and IL-6 are known to be TAM-associated, and both IL-4 and IL-6 are described in the literature as immunosuppressive cytokines that favor a tumor-supportive micromilieu.^25^ When assessing cytokine levels based on WHO grading, we observed significantly increased protein levels in higher-grade tumors for a number of other factors, including the immunosuppressive cytokines IL-8 and IL-10, as well as the angiogenesis-promoting factor VEGF (Suppl. Fig. S3).^15,25–27^

**Figure 2:**
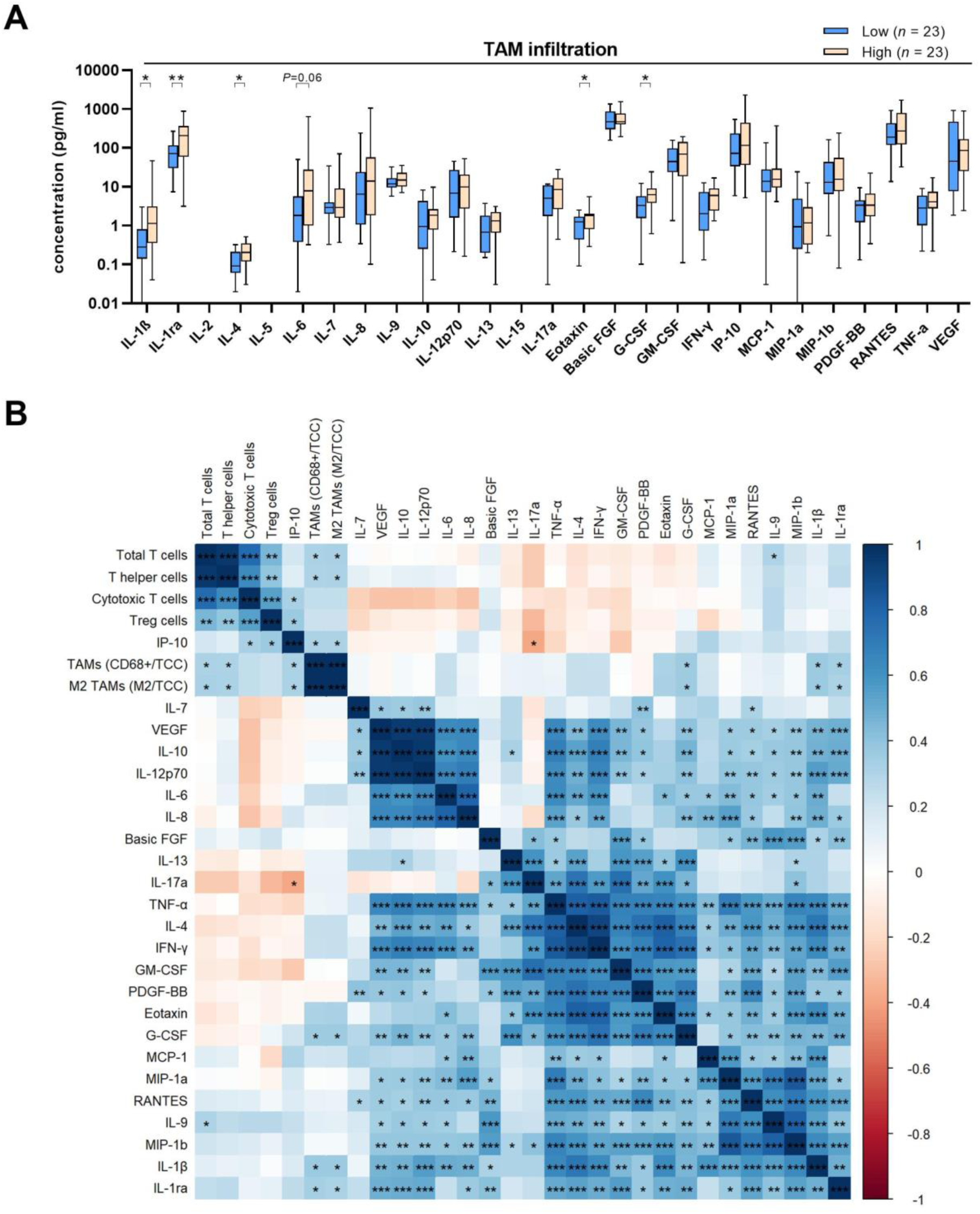
The cyto- and chemokine profile of meningiomas. **(A)** Concentrations of 24 cytokines and chemokines in MGM tissues (n=46) assessed by Luminex analysis comparing TAM low (light blue) and TAM high (orange) infiltration in tumor specimens (median split). **(B)** Correlation matrix of protein concentrations, TAM and TIL (A) using Mann-Whitney-U tests (individually for each analyte) and in (B) using Spearman correlation. Abbreviations: MGM, meningioma; PFS, progression-free survival; TAM, tumor-associated macrophage; TIL, tumor-infiltrating T lymphocyte. Statistical significance: *, *P*<0.05; **, *P*<0.01; ***, *P*<0.001.

To further elucidate the immune network in MGMs, we integrated the protein concentrations and TAM infiltration data from our present study with TIL infiltration data (relative infiltration of CD3^+^ TILs per TCC in MGM tissue in the same patient cohort) from our previous publication^22^ into a correlation matrix for a sub-cohort of *n*=46 cases (Fig. 2B, Suppl. Table S2-3). This resulted in the identification of four distinct clusters, characterized by varying compositions and sizes, and differing degrees of association with the other clusters. The largest cluster contained the pro-inflammatory cytokines IFN-γ and TNF-α as well as the factors Eotaxin, PDGF-BB, G-CSF, and the TAM-associated cytokines GM-CSF and IL-4. Interestingly, especially the pro-inflammatory cytokines IFN-γ and TNF-α strongly correlated with another smaller cluster of several pro-tumoral factors, which are well-described in the literature to create an immunosuppressive niche including IL-6, IL-8, IL-10 and VEGF, and in addition IL12-p70, of which all five factors are also known to be secreted by TAMs in the local TME.^15,25,26^ Furthermore, the last cluster was formed by recruiting chemokines including MCP-1 (CCL2), MIP-1α (CCL3), MIP-1β (CCL4), RANTES (CCL5),^28^ and the cytokines IL-9, IL-1β, IL-1ra. IL-1β acts as a pleiotropic cytokine and has been shown to drive carcinogenesis and metastasis in the tumor context through various mechanisms.^29,30^ Importantly, the balance between IL-1β and its natural antagonist IL-1ra influences the tumor microenvironment’s inflammatory status and impacts TAM polarization and activity.^29,30^ In our analysis, IL-1β and IL-1ra were also significantly correlated with the infiltration of TAMs and M2-TAMs within the first (cellular) cluster.

In summary, quantification of protein levels of 24 immune-related cyto-, chemokines and growth factors revealed a complex TAM-related immunosuppressive micromilieu in MGMs, suggesting TAMs to play an important role therein by secreting a number of immunosuppressive and tumor-supportive cytokines.

### High TAM infiltration outperforms the beneficial prognostic effects of TILs in meningioma

To further elucidate the complex association between macrophages and T cells in MGMs and their distinct impact on patient survival and transcriptional programs, we performed an integrative survival analysis of our TAM infiltration and previously published TIL infiltration data^22^ in the same patient cohort. To this end, the above stated selection criteria (gross total tumor resection (Simpson°I-III), no prior treatment, follow-up >5 years) were applied to the study sample to prevent any survival bias, resulting in a cohort of 94 patients with newly-diagnosed tumors, which were then categorized into four groups according to their combined median TAM and TIL infiltration into (1) low TAM / high TIL, (2) low TAM / low TIL, (3) high TAM / high TIL, (4) high TAM / low TIL infiltration, respectively (Fig. 3A, Suppl. Fig. S4A-C). Significant differences in PFS were observed among the four groups (Fig. 3A; *P*=0.009; log-rank test). Patients with high TIL and low TAM infiltration exhibited superior outcomes, with a median PFS of 146.5 months (*n*=17, light blue). In contrast, the group with low TAM and low TIL numbers (*n*=30, dark blue) exhibited an intermediate survival rate, with a median PFS of 101.9 months. Notably, both groups with high TAM infiltration exhibited the most unfavorable outcomes. The group with high TAM and high TIL infiltration (*n*=30, orange) had a median PFS of 74.9 months and the group with high TAM and low TIL numbers (*n*=17, dark red) had an even worse outcome with a median PFS of 64.5 months. This is of particular interest as the group with high TAM and high TIL infiltration demonstrated comparable high TIL numbers to the group with the most optimal outcome, namely low TAM and high TIL infiltration (Suppl. Fig. S4B). These findings suggest that high TAM infiltration exerts the most dominant negative influence on patient outcome, irrespective of TIL numbers.

**Figure 3:**
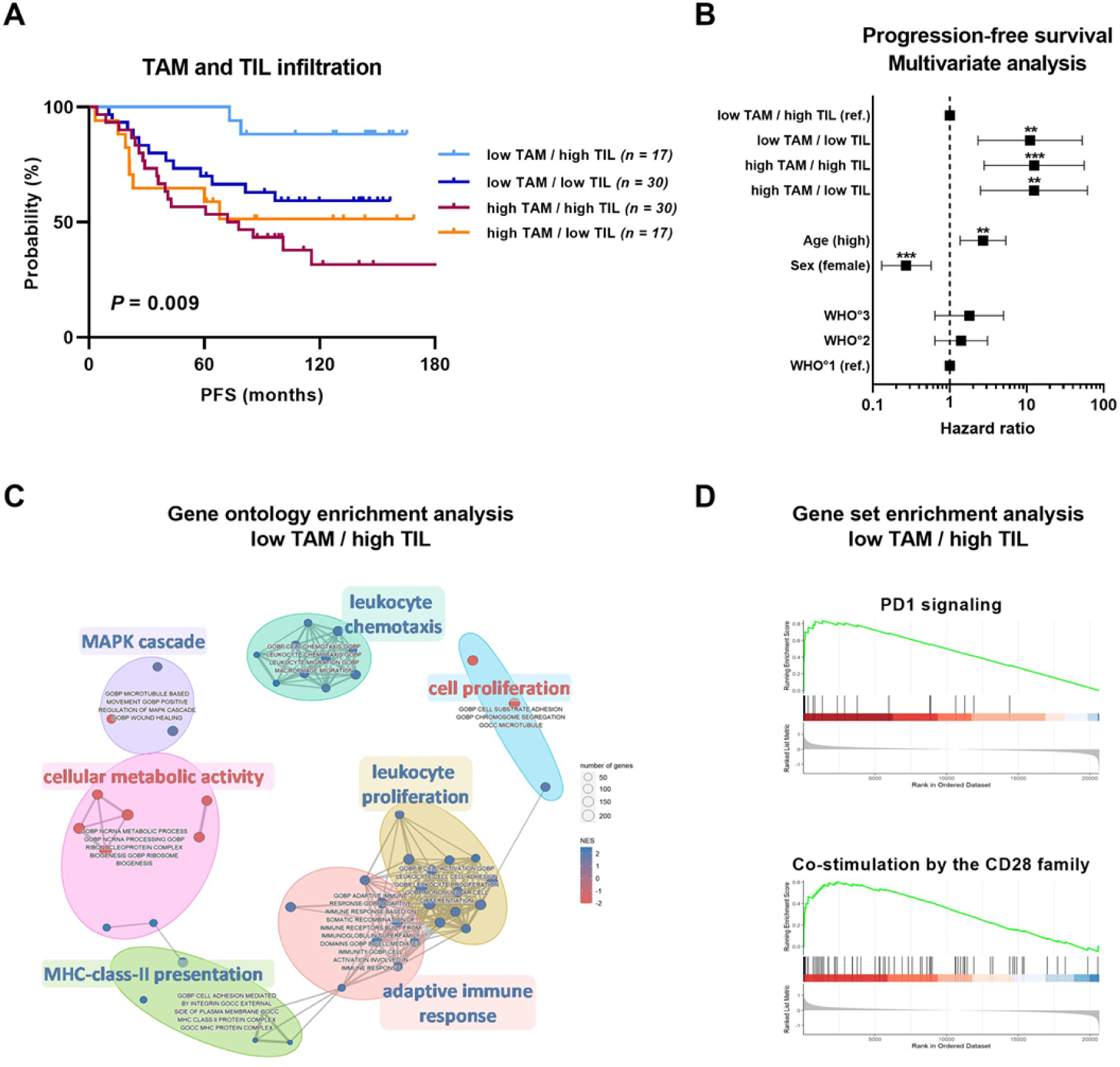
Impact of TAM and TIL infiltration on survival and transcriptional programs in meningiomas. **(A)** Kaplan-Meier plots showing PFS according to combined TAM (CD68^+^/TCC) and TIL (CD3^+^/TCC) infiltration in newly-diagnosed MGMs. The patients were categorized into four groups according to their combined median TAM and TIL infiltration into (1) low TAM / high TIL (light blue), (2) low TAM / low TIL (dark blue), (3) high TAM / high TIL (orange), (4) high TAM / low TIL (dark red) infiltration, respectively. **(B)** Multivariate analysis for PFS including TAM and TIL infiltration grouping, patient age, sex, and WHO grade. Statistical significance was calculated using log-rank test in (A) and Cox proportional hazard model in (B). **(C-D)** Gene expression analysis of microarray data (GSE74385 (*n*=62 cases)^21^ with additional *n*=35 MGM cases) according to the combined median TAM and TIL infiltration in patients with newly-diagnosed and recurrent MGMs. For the analysis, the low TAM / high TIL group was compared to the three remaining groups. **(C)** Gene ontology enrichment analysis in the low TAM / high TIL group. Down-regulated gene sets are depicted in red while up-regulated gene sets are depicted in blue. **(D)** Reactome Pathway Database gene set enrichment analysis in the low TAM / high TIL group showing upregulated PD1 signaling and upregulated co-stimulation by the CD28 family. Abbreviations: MGM, meningioma; PFS, progression-free survival; TAM, tumor-associated macrophage; TCC, total cell count; TIL, tumor-infiltrating T lymphocyte. Statistical significance: *, *P*<0.05; **, *P*<0.01; ***, *P*<0.001.

Most importantly, in a subsequent multivariate analysis, including clinically relevant covariates (age, sex and WHO grade), we were able to show that high TAM and high TIL infiltration (Fig. 3B, Suppl. Table S4; HR 12.48; 95% CI 2.79-55.81; *P*<0.001; Cox PH model) as well as high TAM and low TIL infiltration (Fig. 3B, Suppl. Table S4; HR 12.42; 95% CI 2.50-61.68; *P*=0.002; Cox PH model) are independent prognostic factors for inferior PFS in patients with newly-diagnosed MGMs.

Next, we aimed to explore the impact of the combined TAM and TIL infiltration on the transcriptional programs in MGM tissues by re-analyzing our microarray dataset (GSE74385, *n*=62 cases^21^ with additional *n*=35 MGM cases). To increase statistical power, we used the complete dataset containing both newly-diagnosed and recurrent tumors and performed gene expression analysis contrasting the low TAM and high TIL group with the three remaining groups. Gene ontology (GO) enrichment analysis revealed several significantly shared GO terms in the low TAM and high TIL group, of which GO terms for cellular metabolic activity and cell proliferation were found to be down-regulated (in red; Fig. 3C), whereas GO terms for MAPK cascade, adaptive immune response, MHC class-II presentation, leukocyte chemotaxis and proliferation were found to be up-regulated (in blue; Fig. 3C), indicating an immunologically active microenvironment in tumors with low TAM and high TIL infiltration. In addition, gene set enrichment analysis using the Reactome Pathway Database revealed increased PD1 signaling (normalized enrichment score (NES)=2.398, *P*_adj_<0.0001) and increased T cell signaling represented through co-stimulation by the CD28 family pathway (NES=2.173, *P*_adj_<0.0001) in MGMs with low TAM and high TIL infiltration (Fig. 3D), further supporting an immunostimulatory TME within these tumors.

Taken together, high TAM infiltration - even in cases with high TIL infiltration – was found to be associated with inferior PFS in patients with newly-diagnosed MGMs and was confirmed as an independent prognostic factor in our subsequent multivariate analysis. Further, our gene expression analysis showed that TAM and TIL infiltration are associated with significant changes in the transcriptional profiles of tumors depending on their infiltration status. Thus, our data suggest that TAMs play a vital role in establishing and maintaining the immunosuppressive TME of MGMs and further have a dominant negative impact on patient outcome, which ultimately mitigates the beneficial prognostic effects of TILs in MGM patients.

## DISCUSSION

TAMs represent the main immune cell population in several solid cancer entities, where they are strongly implicated in tumor development and progression,^15,20^ which has also been extensively studied in brain malignancies other than MGMs.^31^ Due to the comparably lower frequency of clinically aggressive MGMs and due to the lack of long-term survival data from patients in most studies, there is still insufficient knowledge about the abundance and functional role of TAMs in this tumor entity, in particular with regard to their influence on tumor behavior and patient outcome. To address these important questions, we broadly investigated TAM and M2-TAM infiltration in a large cohort of 195 clinically well-annotated cases and are the first to evaluate their prognostic value on long-term patient survival (PFS) in MGMs. We discovered a highly heterogeneous but substantial TAM infiltration, which was four times higher than for TILs in the same patient cohort of newly-diagnosed MGMs. ^22^ We further found overall higher numbers of TAMs and pro-tumoral M2-TAMs in clinically aggressive tumors. Importantly, high TAM infiltration turned out as an independent prognostic factor for poor survival outcome in MGM patients dominating over the opposing beneficial prognostic effect of TIL infiltration. Thus, our data provide strong evidence for the immunosuppressive state of TAMs in MGM and their dominant impact on disease progression and survival.

A major strength of our work is the use of a large and clinically well-annotated multicenter study cohort enriched for higher-grade and clinically aggressive MGMs with a distinction between newly-diagnosed (*n*=120) and recurrent (*n*=75) tumors. In addition, our tissue cytometry workflow for assessing TAM numbers and their polarization state in whole-tumor sections rather than the use of small tissue microarrays (TMAs) allows for a highly reliable quantitative analysis. Although, a small number of studies has so far investigated the myeloid cell compartment and reported the presence of TAMs in MGM, most of these studies had major limitations in their study design.^11–14,32–36^ The majority of studies focused on WHO°1 MGMs, while higher-grade and recurrent MGMs were markedly underrepresented, or investigated overall relatively small study cohorts.^13,14,32,33,35,36^ Other studies presented more balanced cohorts with meaningful numbers of higher-grade tumors but primarily focused on expression of the immune checkpoint molecule PD-L1 in MGM tissue,^34,37,38^ or on other cell types, such as MDSCs.^11,12^ In 2019, Proctor and colleagues examined a rather small cohort of *n*=30 MGMs and reported the existence of tumor-supportive M2-TAMs in the immune microenvironment of MGMs.^13^ Further, in 2021, Yeung and colleagues investigated the immunological landscape in a cohort of 201 MGMs using CIBERSORTx, but with a main focus on distinguishing various genetic subtypes in MGM patients. In their *in silico* analysis the authors reported an overall myeloid-enriched immune compartment but did not examine the impact of TAMs on tumor behavior and disease progression.^36^ In another study by Yeung and colleagues, the infiltration of TAMs among other immune cell subsets was examined in a cohort of 73 MGM specimens (*n*=56 WHO°1, *n*=13 WHO°2, *n*=4 WHO°3) using multicolor immunofluorescence stainings on TMA-based small biopsies.^14^ Similar to our findings, their analysis revealed a heterogeneous TAM infiltration in MGMs with the majority of TAMs displaying an immunosuppressive M2 phenotype (CD68^+^ CD163^+^ cells). However, in contrast to our data, the authors reported no significant differences in TAM and M2-TAM infiltration rates across WHO grades, which could be due to differences in study design (staining of comparably small areas vs. here whole-tissue sections) and overall limited number of higher-grade tumors (*n*=13 WHO°2/°3 vs. here *n*=87 WHO°2/°3 newly-diagnosed MGMs). Another drawback of most studies is the limitation to one single technique for examining the TAM compartment,^12–14,33^ whereas in this study, we complemented our tissue cytometry analysis on TAM and M2-TAM infiltration with cytokine analyses in a sub-cohort, and further integrated our previously published datasets on TIL infiltration^22^ as well as gene expression^21^ in MGM tissues, corroborating the predominantly immunosuppressive role of M2-TAMs in this tumor type.

Importantly, a remarkable strength of our analysis is the long-term PFS data with a minimum follow-up of five years, enabling us to investigate longitudinal changes of TAM composition in MGMs as well as their impact on patient survival. Thus, to the best of our knowledge, this is the first comprehensive report demonstrating not only TAM frequencies and their polarization state but also their malignancy- and survival-associated changes in a large study cohort composed of newly-diagnosed as well as recurrent MGMs containing also high frequencies of higher-grade tumors. In our survival analysis, we identified high M2-TAM numbers to be associated with inferior PFS in newly-diagnosed MGM patients and discovered high M2-TAM infiltration as a prognostic factor for poor patient outcome independent of other prognostic confounders, such as age, sex and WHO grade. Moreover, by integrating our previously published data on TIL infiltration^22^ in MGM in the same patient cohort, we analyzed the combined TAM and TIL infiltration and identified high TAM infiltration as the dominant negative prognostic factor on patient outcome in MGM tissue, highlighting the importance of TAMs on tumor behavior and disease progression. Based on our findings, we promote TAMs and M2-TAMs as potential therapeutic targets for immunotherapeutic approaches in MGM patients.

To date, clinical trials have primarily focused on T cell-based immune checkpoint inhibition, despite substantial evidence that MGMs largely do not meet essential prerequisites facilitating the clinical success of ICB therapy, which are in general (1) high tumor mutational burden, (2) high infiltration of T cells, and (3) high expression of ICB targets.^10,39^ Accordingly, it is no surprise that results from clinical trials targeting the PD1/PD-L1 axis or CTLA4, either as mono- or combination therapy, have been fairly disappointing in MGM patients.^10,40,41^ Fortunately, novel immunotherapeutic strategies have entered pre-clinical and clinical testing in recent years and have focused among others on targeting immunosuppressive macrophages at the tumor site.^20,42,43^ TAMs as treatment targets have attracted great interest, especially in brain malignancies, such as gliomas, which are also characterized by a low tumor mutational burden, lower infiltration of T cells but higher numbers of TAMs.^24,31,44^ In 2021, Yeung and colleagues were the first to investigate a macrophage-targeting approach in an immune-competent syngeneic mouse model of MGM and reported anti-CSF1/CSF1R immunotherapy to be efficacious at inhibiting tumor growth,^36^ which has previously been reported to elicit anti-tumoral responses in other brain tumor models as well.^45–47^ As a consequence, we further encourage to rethink immunotherapeutic approaches for MGM patients.^41^ Finally, with regard to our survival analysis in recurrent MGMs, where the presence of TAMs had no further impact on survival, we additionally propose to consider the timing of immunotherapeutic approaches in MGM patients. In the past, clinical trials have primarily enrolled patients in an advanced disease stage, with clinically aggressive tumors that have been heavily pre-treated and therefore likely have a highly immunosuppressive microenvironment, in which immunotherapeutic interventions, regardless of the target, may ultimately fail to induce clinically meaningful responses. Therefore, future clinical trials of immunotherapy targeting macrophages in MGM should be favorably given in the primary setting rather than at recurrence and could be eventually combined with radiotherapy or other T cell-based immunotherapies to improve overall patient outcome.

In summary, we performed a comprehensive integrative analysis of TAM and TIL infiltration, immune-related cytokines and gene expression in a clinically well-annotated large study cohort (*n*=195) encompassing both newly-diagnosed and recurrent MGMs as well as substantial numbers of higher-grade tumors. Thereby, we found high M2-TAM infiltration to be associated with inferior PFS in MGM patients, and importantly, identified high TAM infiltration as an independent prognostic factor for poor patient outcome, dominating the opposing beneficial prognostic effect of TILs by creating an immunosuppressive niche. Based on our findings, we strongly suggest M2-TAMs as attractive treatment targets for immunotherapeutic clinical trials in MGM patients, hopefully leading to novel therapeutic approaches in the near future.

## Authorship

**Conception and design:** C.L., F.L., R.W., C.R., C.H.M.

**Development of methodology:** F.L., R.W., C.R., M.B., N.G., C.H.M.

**Acquisition of data:** C.L., F.L., R.W., G.J., C.R., M.B., K.L., A.F.K., N.G., M.L., R.K., C.S., M.W., F.S., S.K., A.U., M.S., A.v.D., C.H.M.

**Analysis and interpretation of data:** C.L., F.L., R.W., G.J., C.R., M.B., K.L., M.L., R.K., M.S., C.H.M.

**Writing – original draft:** C.L., F.L., R.W., G.J., C.H.M.

**Writing - review, and/or revision of the manuscript:** all authors.

**Administrative, technical, or material support:** C.L., F.L., R.W., G.J., C.R., M.B., K.L., A.F.K., N.G., M.L., R.K., C.S., M.W., F.S., S.K., A.U., M.S., A.v.D., C.H.M.

**Study supervision:** R.W., C.R., C.H.M.

## Data availability

All data are being securely held within the Division of Experimental Neurosurgery, Department of Neurosurgery at the University Hospital of Heidelberg, Germany. All data have been systematically cataloged and are readily available. Microarray dataset GSE74385 is available online.

## Acknowledgments

We thank the German Cancer Research Center (DKFZ, Heidelberg, Germany) Omics IT Data Management Core Facility (ODCF) and the Genomics Core Facilities for providing excellent technical support.

## SUPPLEMENTARY FIGURES

**Supplementary Figure S1:**
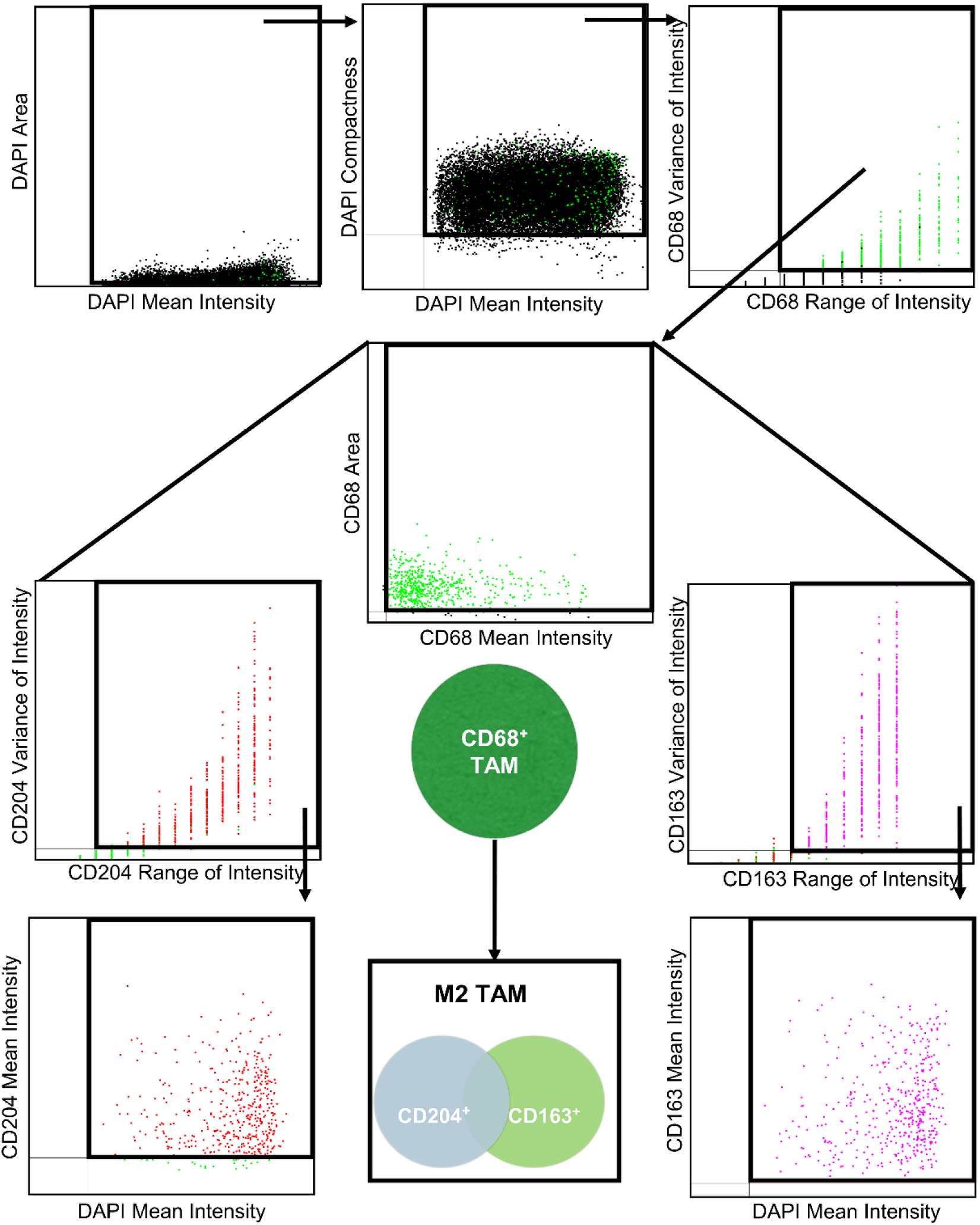
Workflow for tissue cytometry-based analysis. Automated cell detection was based on DAPI. Irregular and extreme small and large DAPI were excluded at first. Subsequent detection of marker CD68 was conducted within the gate of the selected DAPI. Before the assessment to the three macrophage markers, unspecific homogenous background and dotted artifacts were filtered by the setting of area as well as general TAM. Analysis of CD204 and CD163 were performed within the gate of CD68 positive cells. CD68^+^ cells with staining for CD204^+^ and/or CD163^+^ were recognized as M2-TAM. Abbreviations: TAM, tumor-associated macrophage.

**Supplementary Figure S2:**
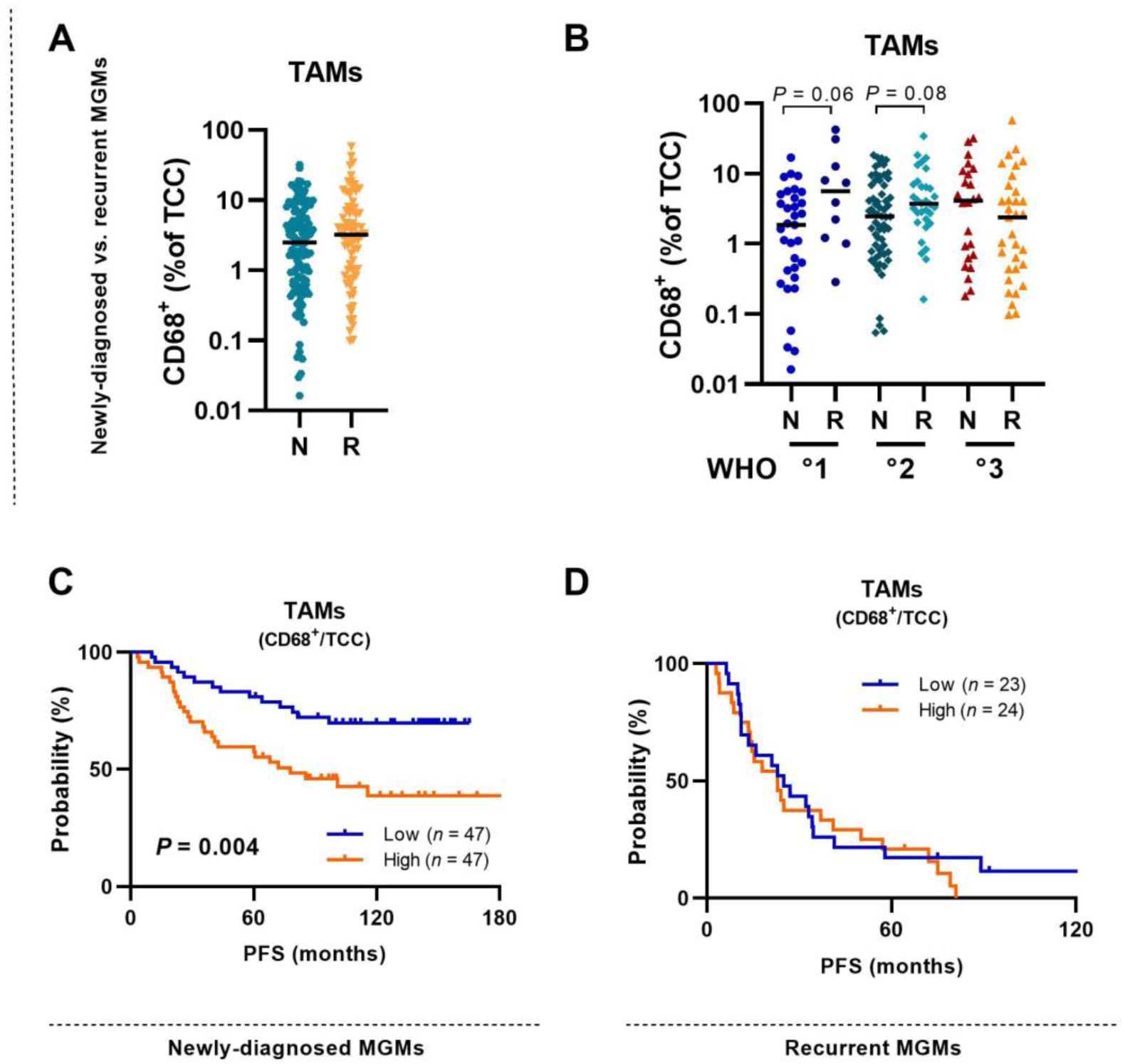
TAM infiltration and impact on survival in newly-diagnosed and recurrent meningiomas. **(A)** TAM infiltration (CD68^+^/TCC) in newly-diagnosed (N) and recurrent (R) MGMs. **(B)** TAM infiltration across WHO grades in newly-diagnosed (N) and recurrent (R) MGMs. **(C-D)** Kaplan-Meier plot for PFS based on high (orange curve) and low (blue curve) TAM infiltration in **(C)** newly-diagnosed and **(D)** recurrent MGMs Statistical significance was calculated using Mann-Whitney-U test in (A-B), and log-rank test in (C-D). Abbreviations: MGM, meningioma; N, newly-diagnosed; PFS, progression-free survival; R, recurrent; TAM, tumor-associated macrophage; TCC, total cell count. Statistical significance: *, *P*<0.05; **, *P*<0.01.

**Supplementary Figure S3:**
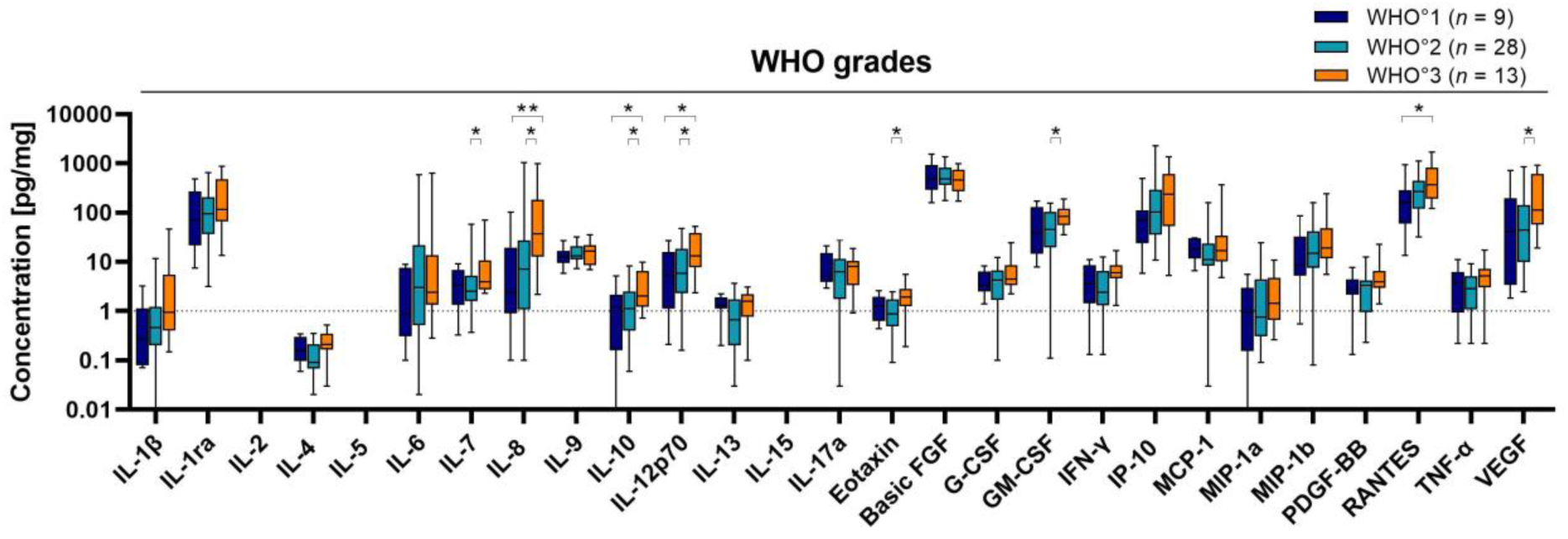
The cyto- and chemokine profile of meningiomas across WHO grades. Concentrations of 24 cytokines and chemokines in MGM tissues (n=46) assessed by Luminex analysis comparing WHO grades. Statistical significance was calculated using Mann-Whitney-U test. Abbreviations: MGM, meningioma. Statistical significance: *, *P*<0.05; **, *P*<0.01.

**Supplementary Figure S4:**
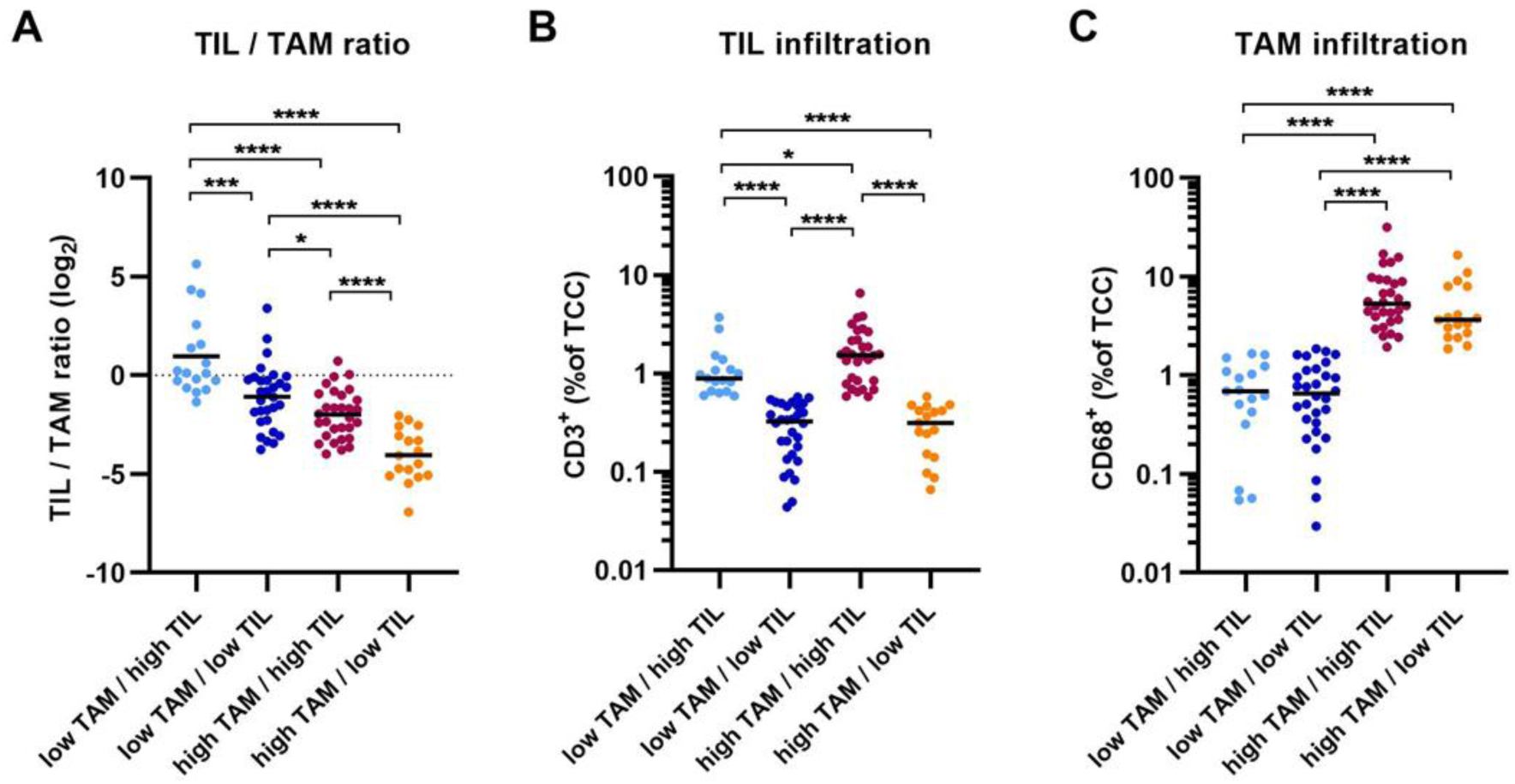
TAM and TIL infiltration in newly-diagnosed meningiomas. **(A-C)** TAM and TIL infiltration in newly-diagnosed MGMs showing **(A)** TIL / TAM ratio, **(B)** TIL infiltration (CD3^+^/TCC) and **(C)** TAM infiltration (CD68^+^/TCC) across the four specified groups of patients according to their median TAM and TIL infiltration: (1) low TAM / high TIL (light blue), (2) low TAM / low TIL (dark blue), (3) high TAM / high TIL (orange), (4) high TAM / low TIL (dark red) infiltration, respectively. Lines show the mean in (A) and the median in (B-C). Statistical significance was calculated by Student’s unpaired t test in (A) and by Mann-Whitney-U test in (B-C). Abbreviations: TAM, tumor-associated macrophage; TIL, tumor-infiltrating T lymphocyte. Statistical significance: *, *P*<0.05; **, *P*<0.01; ***, *P*<0.001; ****, *P*<0.0001.

## SUPPLEMENTARY TABLES

**Supplementary Table S1:** Impact of M2-TAM infiltration on patient survival. Multivariate analysis (Cox proportional hazard model).

| PROGRESSION-FREE SURVIVAL |  |  |  |  |  |
| --- | --- | --- | --- | --- | --- |
|  |  | n | HR | 95%-CI | P-value |
| M2-TAM INFILTRATION | low | 47 | 1.00 |  |  |
|  | high | 47 | 2.11 | 1.11-4.01 | **0.023\*** |
| AGE | low | 47 | 1.00 |  |  |
|  | high | 47 | 2.06 | 1.07-3.94 | **0.030\*** |
| SEX | male | 36 | 1.00 |  |  |
|  | female | 58 | 0.40 | 0.20-0.78 | **0.007\*\*** |
| WHO° | 1 | 27 | 1.00 |  |  |
|  | 2 | 54 | 1.04 | 0.48-2.27 | 0.914 |
|  | 3 | 13 | 1.62 | 0.60-4.35 | 0.336 |

Results of the multivariate analysis for the progression-free survival of newly-diagnosed meningioma cases calculated using Cox proportional hazard model. Abbreviations: 95%-CI, lower and upper border of 95% confidence interval; HR, hazard ratio; n, number; TAM, tumor-associated macrophage.

**Supplementary Table S2:** Correlation coefficients derived from correlation matrix of protein concentrations, TAM and TIL infiltration numbers ordered by Spearman correlation. Color code of cells derived from correlation matrix in Figure 2D.

|  | Total T cells | T helper cells | Cytotoxic T cells | Treg cells | IP-10 | TAMs (CD68+/TCC) | M2 TAMs (M2/TCC) | IL-7 | VEGF | IL-10 | IL-12p70 | IL-6 | IL-8 | Basic FGF | IL-13 | IL-17a | TNF-α | IL-4 | IFN-γ | GM-CSF | PDGF-BB | Eotaxin | G-CSF | MCP-1 | MIP-1a | RANTES | IL-9 | MIP-1b | IL-1β | IL-1ra |
| --- | --- | --- | --- | --- | --- | --- | --- | --- | --- | --- | --- | --- | --- | --- | --- | --- | --- | --- | --- | --- | --- | --- | --- | --- | --- | --- | --- | --- | --- | --- |
| Total T cells | 1.0000 | 0.9609 | 0.7716 | 0.4579 | 0.1899 | 0.3168 | 0.3224 | 0.0270 | -0.0221 | -0.0001 | 0.0154 | -0.0286 | -0.0717 | 0.1153 | -0.1016 | -0.2540 | -0.0463 | -0.1582 | -0.1043 | -0.1142 | -0.0749 | -0.1305 | -0.0880 | 0.1314 | 0.0282 | 0.1577 | 0.3005 | 0.2023 | 0.1256 | 0.2460 |
| T helper cells | 0.9609 | 1.0000 | 0.6350 | 0.3919 | 0.1338 | 0.3043 | 0.3140 | 0.1175 | 0.0694 | 0.1002 | 0.1161 | 0.0806 | 0.0104 | 0.0542 | -0.1151 | -0.2556 | -0.0072 | -0.1384 | -0.0483 | -0.1081 | -0.0459 | -0.0937 | -0.0667 | 0.1105 | 0.0184 | 0.1315 | 0.2522 | 0.1773 | 0.0972 | 0.2681 |
| Cytotoxic T cells | 0.7716 | 0.6350 | 1.0000 | 0.5462 | 0.3233 | 0.2426 | 0.2444 | -0.2369 | -0.2890 | -0.2803 | -0.2794 | -0.2302 | -0.2885 | 0.1315 | -0.0217 | -0.1410 | -0.1723 | -0.0990 | -0.1585 | -0.1879 | -0.0304 | -0.0233 | -0.0420 | -0.0126 | -0.0192 | 0.1241 | 0.2791 | 0.1521 | 0.1032 | 0.0826 |
| Treg cells | 0.4579 | 0.3919 | 0.5462 | 1.0000 | 0.3614 | 0.2197 | 0.2178 | -0.2041 | -0.1110 | -0.0821 | -0.0876 | -0.0591 | -0.1381 | 0.2093 | -0.1058 | -0.3282 | -0.1502 | -0.0736 | -0.0472 | -0.1770 | -0.0627 | -0.0679 | 0.0633 | -0.1964 | -0.0592 | 0.0571 | 0.2715 | 0.0999 | 0.0249 | 0.1077 |
| IP-10 | 0.1899 | 0.1338 | 0.3233 | 0.3614 | 1.0000 | 0.3182 | 0.3110 | -0.0722 | -0.0576 | -0.0573 | -0.0573 | 0.0357 | 0.1832 | 0.0448 | -0.0809 | -0.3801 | -0.2018 | -0.0559 | -0.0645 | -0.2616 | -0.0315 | -0.0390 | 0.2078 | 0.3003 | 0.1746 | 0.1405 | 0.2350 | 0.1916 | 0.1533 | 0.2522 |
| TAMs (CD68+/TCC) | 0.3168 | 0.3043 | 0.2426 | 0.2197 | 0.3182 | 1.0000 | 0.9965 | 0.0502 | -0.0354 | 0.0459 | 0.0144 | 0.2248 | 0.0757 | 0.0121 | 0.0831 | 0.0972 | 0.0782 | 0.2370 | 0.2226 | 0.0134 | -0.0491 | 0.3115 | 0.3587 | 0.1965 | 0.0118 | 0.0960 | 0.1240 | 0.0483 | 0.3607 | 0.3377 |
| M2 TAMs (M2/TCC) | 0.3224 | 0.3140 | 0.2444 | 0.2178 | 0.3110 | 0.9965 | 1.0000 | 0.0560 | -0.0359 | 0.0478 | 0.0163 | 0.2367 | 0.0806 | 0.0101 | 0.0876 | 0.0977 | 0.0847 | 0.2312 | 0.2350 | 0.0088 | -0.0415 | 0.3212 | 0.3577 | 0.2100 | 0.0289 | 0.0822 | 0.1188 | 0.0466 | 0.3676 | 0.3180 |
| IL-7 | 0.0270 | 0.1175 | -0.2369 | -0.2041 | -0.0722 | 0.0502 | 0.0560 | 1.0000 | 0.3722 | 0.3538 | 0.3813 | 0.1335 | 0.1034 | 0.1192 | 0.2851 | -0.1317 | 0.1811 | 0.0772 | 0.1456 | 0.2299 | 0.4071 | 0.0515 | 0.2811 | 0.3134 | 0.1228 | 0.3144 | 0.1759 | 0.1522 | 0.0789 | 0.2424 |
| VEGF | -0.0221 | 0.0694 | -0.2890 | -0.1110 | -0.0576 | -0.0354 | -0.0359 | 0.3722 | 1.0000 | 0.9583 | 0.9583 | 0.6307 | 0.6464 | 0.1888 | 0.2892 | -0.0951 | 0.6474 | 0.4488 | 0.6224 | 0.4529 | 0.3867 | 0.2634 | 0.4035 | 0.2860 | 0.3598 | 0.3519 | 0.3390 | 0.3877 | 0.4625 | 0.4812 |
| IL-10 | -0.0001 | 0.1002 | -0.2803 | -0.0821 | -0.0573 | 0.0459 | 0.0478 | 0.3538 | 0.9583 | 1.0000 | 0.9676 | 0.5978 | 0.6418 | 0.1136 | 0.3232 | -0.0468 | 0.6858 | 0.5319 | 0.6879 | 0.4552 | 0.3926 | 0.3265 | 0.4602 | 0.2603 | 0.3604 | 0.3339 | 0.3702 | 0.3894 | 0.4783 | 0.4934 |
| IL-12p70 | 0.0154 | 0.1161 | -0.2794 | -0.0876 | -0.0573 | 0.0144 | 0.0163 | 0.3813 | 0.9583 | 0.9676 | 1.0000 | 0.6553 | 0.6930 | 0.0958 | 0.2282 | -0.0602 | 0.6283 | 0.4823 | 0.6570 | 0.4048 | 0.3257 | 0.3333 | 0.4098 | 0.2758 | 0.3933 | 0.3849 | 0.3533 | 0.3827 | 0.5526 | 0.5038 |
| IL-6 | -0.0286 | 0.0806 | -0.2302 | -0.0591 | 0.0357 | 0.2248 | 0.2367 | 0.1335 | 0.6307 | 0.5978 | 0.6553 | 1.0000 | 0.8064 | 0.0138 | 0.1517 | 0.1377 | 0.6354 | 0.5128 | 0.5988 | 0.1822 | 0.2694 | 0.4219 | 0.3794 | 0.3773 | 0.4332 | 0.3862 | 0.3466 | 0.3895 | 0.4986 | 0.2324 |
| IL-8 | -0.0717 | 0.0104 | -0.2885 | -0.1381 | 0.1832 | 0.0757 | 0.0806 | 0.1034 | 0.6464 | 0.6418 | 0.6930 | 0.8064 | 1.0000 | 0.0309 | 0.0841 | -0.1757 | 0.5357 | 0.3725 | 0.5133 | 0.2727 | 0.2162 | 0.3380 | 0.4307 | 0.4664 | 0.5436 | 0.3775 | 0.2401 | 0.3912 | 0.4896 | 0.3652 |
| Basic FGF | 0.1153 | 0.0542 | 0.1315 | 0.2093 | 0.0448 | 0.0121 | 0.0101 | 0.1192 | 0.1888 | 0.1136 | 0.0958 | 0.0138 | 0.0309 | 1.0000 | 0.1879 | 0.4204 | 0.3611 | 0.2527 | 0.2834 | 0.5618 | 0.3875 | 0.2157 | 0.2287 | 0.3117 | 0.3624 | 0.4670 | 0.5743 | 0.5525 | 0.3127 | 0.4130 |
| IL-13 | -0.1016 | -0.1151 | -0.0217 | -0.1058 | -0.0809 | 0.0831 | 0.0876 | 0.2851 | 0.2892 | 0.3232 | 0.2282 | 0.1517 | 0.0841 | 0.1879 | 1.0000 | 0.5956 | 0.3637 | 0.5341 | 0.3319 | 0.5839 | 0.5473 | 0.3992 | 0.6067 | 0.1532 | 0.1186 | 0.1566 | 0.2242 | 0.3083 | 0.3038 | 0.1823 |
| IL-17a | -0.2540 | -0.2556 | -0.1410 | -0.3282 | -0.3801 | 0.0972 | 0.0977 | -0.1317 | -0.0951 | -0.0468 | -0.0602 | 0.1377 | -0.1757 | 0.4204 | 0.5956 | 1.0000 | 0.5207 | 0.7341 | 0.5672 | 0.7585 | 0.5140 | 0.6084 | 0.4577 | 0.1963 | 0.1594 | 0.1960 | 0.2157 | 0.4015 | 0.3042 | 0.2365 |
| TNF-α | -0.0463 | -0.0072 | -0.1723 | -0.1502 | -0.2018 | 0.0782 | 0.0847 | 0.1811 | 0.6474 | 0.6858 | 0.6283 | 0.6354 | 0.5357 | 0.3611 | 0.3637 | 0.5207 | 1.0000 | 0.8110 | 0.8366 | 0.6730 | 0.6754 | 0.6691 | 0.5903 | 0.4378 | 0.6959 | 0.5749 | 0.5096 | 0.6644 | 0.6405 | 0.5076 |
| IL-4 | -0.1582 | -0.1384 | -0.0990 | -0.0736 | -0.0559 | 0.2370 | 0.2312 | 0.0772 | 0.4488 | 0.5319 | 0.4823 | 0.5128 | 0.3725 | 0.2527 | 0.5341 | 0.7341 | 0.8110 | 1.0000 | 0.9122 | 0.6668 | 0.7492 | 0.8189 | 0.6675 | 0.3512 | 0.4933 | 0.5821 | 0.4817 | 0.5943 | 0.7291 | 0.5345 |
| IFN-γ | -0.1043 | -0.0483 | -0.1585 | -0.0472 | -0.0645 | 0.2226 | 0.2350 | 0.1456 | 0.6224 | 0.6879 | 0.6570 | 0.5988 | 0.5133 | 0.2834 | 0.3319 | 0.5672 | 0.8366 | 0.9122 | 1.0000 | 0.6198 | 0.6745 | 0.8022 | 0.7423 | 0.3746 | 0.4126 | 0.5166 | 0.4416 | 0.5288 | 0.6302 | 0.4985 |
| GM-CSF | -0.1142 | -0.1081 | -0.1879 | -0.1770 | -0.2616 | 0.0134 | 0.0088 | 0.2299 | 0.4529 | 0.4552 | 0.4048 | 0.1822 | 0.2727 | 0.5618 | 0.5839 | 0.7585 | 0.6730 | 0.6668 | 0.6198 | 1.0000 | 0.6665 | 0.5767 | 0.5631 | 0.2041 | 0.3118 | 0.5010 | 0.3073 | 0.5381 | 0.4458 | 0.5544 |
| PDGF-BB | -0.0749 | -0.0459 | -0.0304 | -0.0627 | -0.0315 | -0.0491 | -0.0415 | 0.4071 | 0.3867 | 0.3926 | 0.3257 | 0.2694 | 0.2162 | 0.3875 | 0.5473 | 0.5140 | 0.6754 | 0.7492 | 0.6745 | 0.6665 | 1.0000 | 0.5925 | 0.6940 | 0.2347 | 0.4484 | 0.6579 | 0.3837 | 0.5383 | 0.3728 | 0.3781 |
| Eotaxin | -0.1305 | -0.0937 | -0.0233 | -0.0679 | -0.0390 | 0.3115 | 0.3212 | 0.0515 | 0.2634 | 0.3265 | 0.3333 | 0.4219 | 0.3380 | 0.2157 | 0.3992 | 0.6084 | 0.6691 | 0.8189 | 0.8022 | 0.5767 | 0.5925 | 1.0000 | 0.6494 | 0.3741 | 0.3843 | 0.4765 | 0.3564 | 0.5612 | 0.6307 | 0.4862 |
| G-CSF | -0.0880 | -0.0667 | -0.0420 | 0.0633 | 0.2078 | 0.3587 | 0.3577 | 0.2811 | 0.4035 | 0.4602 | 0.4098 | 0.3794 | 0.4307 | 0.2287 | 0.6067 | 0.4577 | 0.5903 | 0.6675 | 0.7423 | 0.5631 | 0.6940 | 0.6494 | 1.0000 | 0.2335 | 0.3805 | 0.4210 | 0.4137 | 0.4669 | 0.4583 | 0.4931 |
| MCP-1 | 0.1314 | 0.1105 | -0.0126 | -0.1964 | 0.3003 | 0.1965 | 0.2100 | 0.3134 | 0.2860 | 0.2603 | 0.2758 | 0.3773 | 0.4664 | 0.3117 | 0.1532 | 0.1963 | 0.4378 | 0.3512 | 0.3746 | 0.2041 | 0.2347 | 0.3741 | 0.2335 | 1.0000 | 0.5794 | 0.3763 | 0.3598 | 0.4362 | 0.5808 | 0.2330 |
| MIP-1a | 0.0282 | 0.0184 | -0.0192 | -0.0592 | 0.1746 | 0.0118 | 0.0289 | 0.1228 | 0.3598 | 0.3604 | 0.3933 | 0.4332 | 0.5436 | 0.3624 | 0.1186 | 0.1594 | 0.6959 | 0.4933 | 0.4126 | 0.3118 | 0.4484 | 0.3843 | 0.3805 | 0.5794 | 1.0000 | 0.5616 | 0.7020 | 0.8472 | 0.5237 | 0.3673 |
| RANTES | 0.1577 | 0.1315 | 0.1241 | 0.0571 | 0.1405 | 0.0960 | 0.0822 | 0.3144 | 0.3519 | 0.3339 | 0.3849 | 0.3862 | 0.3775 | 0.4670 | 0.1566 | 0.1960 | 0.5749 | 0.5821 | 0.5166 | 0.5010 | 0.6579 | 0.4765 | 0.4210 | 0.3763 | 0.5616 | 1.0000 | 0.6199 | 0.7101 | 0.5962 | 0.5251 |
| IL-9 | 0.3005 | 0.2522 | 0.2791 | 0.2715 | 0.2350 | 0.1240 | 0.1188 | 0.1759 | 0.3390 | 0.3702 | 0.3533 | 0.3466 | 0.2401 | 0.5743 | 0.2242 | 0.2157 | 0.5096 | 0.4817 | 0.4416 | 0.3073 | 0.3837 | 0.3564 | 0.4137 | 0.3598 | 0.7020 | 0.6199 | 1.0000 | 0.8113 | 0.5496 | 0.4094 |
| MIP-1b | 0.2023 | 0.1773 | 0.1521 | 0.0999 | 0.1916 | 0.0483 | 0.0466 | 0.1522 | 0.3877 | 0.3894 | 0.3827 | 0.3895 | 0.3912 | 0.5525 | 0.3083 | 0.4015 | 0.6644 | 0.5943 | 0.5288 | 0.5381 | 0.5383 | 0.5612 | 0.4669 | 0.4362 | 0.8472 | 0.7101 | 0.8113 | 1.0000 | 0.6368 | 0.5341 |
| IL-1β | 0.1256 | 0.0972 | 0.1032 | 0.0249 | 0.1533 | 0.3607 | 0.3676 | 0.0789 | 0.4625 | 0.4783 | 0.5526 | 0.4986 | 0.4896 | 0.3127 | 0.3038 | 0.3042 | 0.6405 | 0.7291 | 0.6302 | 0.4458 | 0.3728 | 0.6307 | 0.4583 | 0.5808 | 0.5237 | 0.5962 | 0.5496 | 0.6368 | 1.0000 | 0.4878 |
| IL-1ra | 0.2460 | 0.2681 | 0.0826 | 0.1077 | 0.2522 | 0.3377 | 0.3180 | 0.2424 | 0.4812 | 0.4934 | 0.5038 | 0.2324 | 0.3652 | 0.4130 | 0.1823 | 0.2365 | 0.5076 | 0.5345 | 0.4985 | 0.5544 | 0.3781 | 0.4862 | 0.4931 | 0.2330 | 0.3673 | 0.5251 | 0.4094 | 0.5341 | 0.4878 | 1.0000 |

**Supplementary Table S3:**
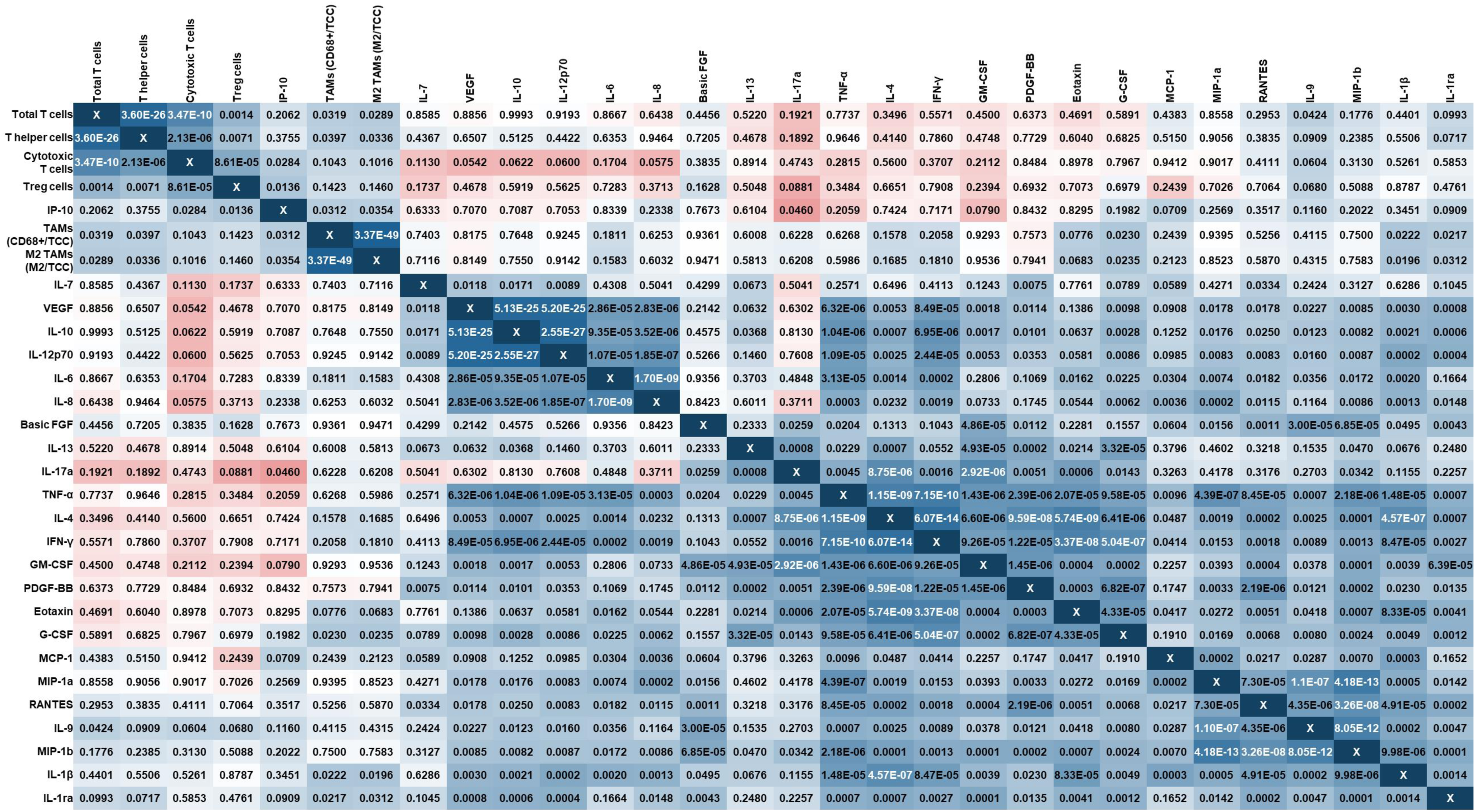
*P*-values derived from correlation matrix of protein concentrations, TAM and TIL infiltration numbers ordered by Spearman correlation. Color code of cells derived from correlation matrix in Figure 2D.

**Supplementary Table S4:**
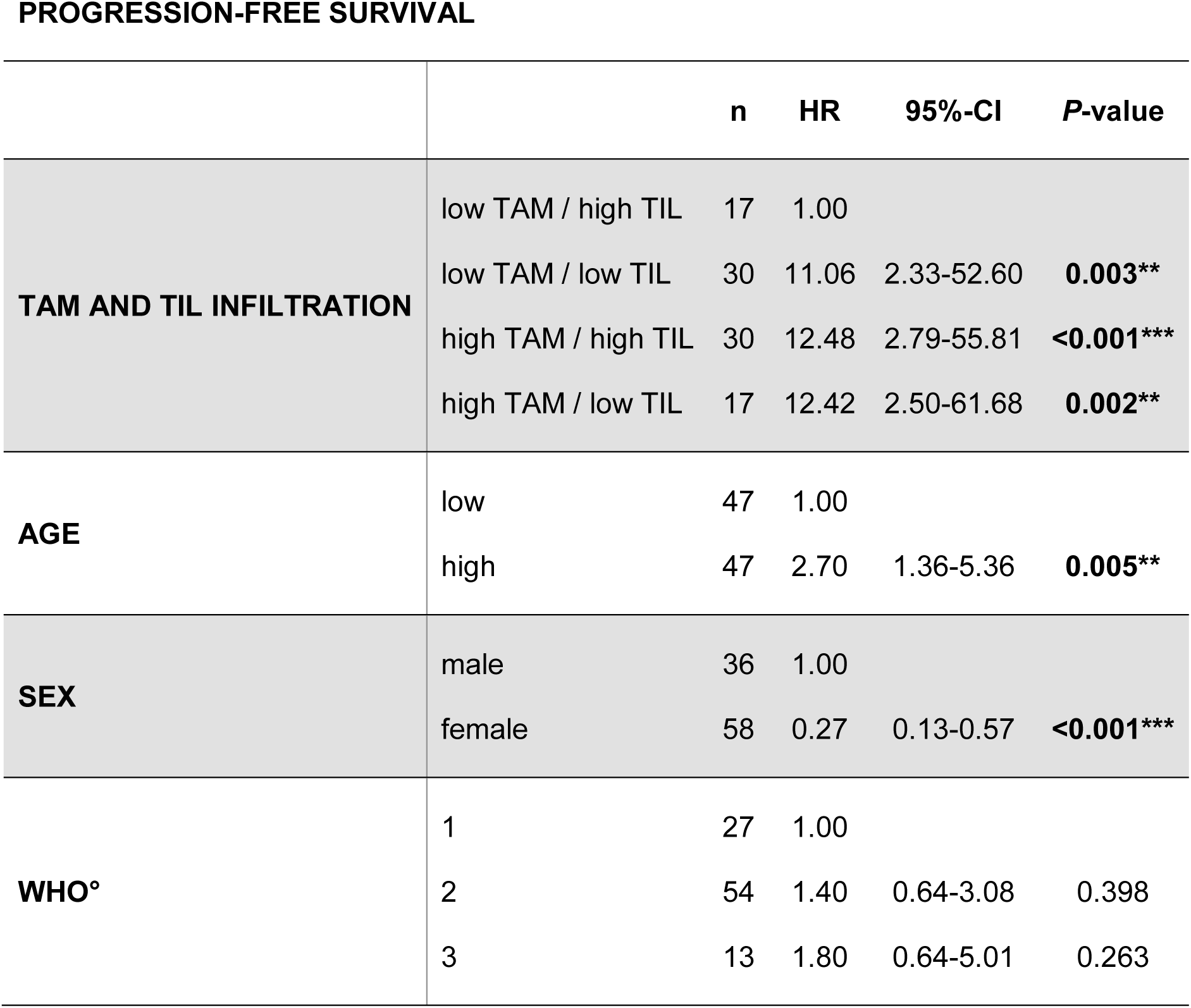
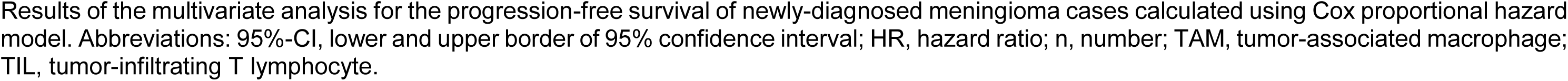
Impact of TAM and TIL infiltration on patient survival. Multivariate analysis (Cox proportional hazard model).

**PROGRESSION-FREE SURVIVAL**
|  |  | **n** | **HR** | **95%-CI** | **P-value** |
| --- | --- | --- | --- | --- | --- |
| **TAM AND TIL INFILTRATION** | low TAM / high TIL | 17 | 1.00 |  |  |
|  | low TAM / low TIL | 30 | 11.06 | 2.33-52.60 | **0.003\*\*** |
|  | high TAM / high TIL | 30 | 12.48 | 2.79-55.81 | **<0.001\*\*\*** |
|  | high TAM / low TIL | 17 | 12.42 | 2.50-61.68 | **0.002\*\*** |
| **AGE** | low | 47 | 1.00 |  |  |
|  | high | 47 | 2.70 | 1.36-5.36 | **0.005\*\*** |
| **SEX** | male | 36 | 1.00 |  |  |
|  | female | 58 | 0.27 | 0.13-0.57 | **<0.001\*\*\*** |
| **WHO°** | 1 | 27 | 1.00 |  |  |
|  | 2 | 54 | 1.40 | 0.64-3.08 | 0.398 |
|  | 3 | 13 | 1.80 | 0.64-5.01 | 0.263 |

Results of the multivariate analysis for the progression-free survival of newly-diagnosed meningioma cases calculated using Cox proportional hazard model. Abbreviations: 95%-CI, lower and upper border of 95% confidence interval; HR, hazard ratio; n, number; TAM, tumor-associated macrophage; TIL, tumor-infiltrating T lymphocyte.

